# Information-Efficient, Off-Center Sampling Results in Improved Precision in 3D Single Particle Tracking Microscopy

**DOI:** 10.1101/2021.03.21.436327

**Authors:** Chen Zhang, Kevin Welsher

## Abstract

In this work, we present a 3D single-particle tracking system that can apply tailored sampling patterns to selectively extract photons that yield the most information for particle localization. We demonstrate that off-center sampling at locations predicted by Fisher information utilizes photons most efficiently. When performing localization in a single dimension, optimized off-center sampling patterns gave doubled precision compared to uniform sampling. A ~20% increase in precision compared to uniform sampling can be achieved when a similar off-center pattern is used in 3D localization. Here we systematically investigated the photon efficiency of different emission patterns in a diffraction-limited system and achieved higher precision than uniform sampling. The ability to maximize information from the limited number of photons demonstrated here is critical for particle tracking applications in biological samples, where photons may be limited.

## Introduction

Single-particle tracking (SPT)[1] has led to numerous advances in unveiling sophisticated intracellular biophysical events, including diffusion of membrane proteins [2], transportation of intracellular vesicles [3], and viral internalization events [4, 5]. Despite the tremendous progress made in the field, there are limits to what can be gleaned from single-particle trajectories by the intrinsic localization precision. In a conventional microscope, this limit is given by 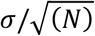 where σ is the size of the microscope’s point-spread function (PSF) and *N* is the number of photons collected [6, 7]. The diffraction limit dictates the size of the PSF, so increasing the number of photons collected per localization is typically the only method available for increasing prevision. While the use of artifitial particles [8] can produce a higher flux of photons and improve localization precision, conventional organic fluorophores and fluorescent proteins remain essential in biophysical studies. These probes can only yield a finite number of photons before undergoing irreversible photobleaching, so it is crucial to maximize the information available from this limited number of photons. In a typical particle tracking experiment, the particle is uniformly illuminated as, typically, the emitter’s position is not known at the outset of the experiment. An under-explored avenue for increasing precision is adjusting the excitation pattern around the emitter to get beyond the localization limit described above. This type of advance is only possible if the particle position is known *a priori*, at least to some degree of certainty. Recent efforts have focused on improving localization precision through non-uniform illumination in the context of a super-resolution technique [9]. Gallatin et al. proposed a globally optimized strategy in which the particle should be sampled at the maximum of the first-order derivative of the square-root of the intensity [10]. This theory indicated that the optimized sampling pattern is non-uniform, and the particle position should not be directly sampled but did not provide experimental support. Both of these works suggest that non-uniform illumination has promised for improved localization with a limited number of photons.

This study builds upon 3D single-molecule active real-time tracking microscopy (3D-SMART)[11], a previously introduced real-time 3D single-particle tracking (RT-3D-SPT) system [12], by utilizing the emerging concept of non-uniform illumination to improve localization precision significantly. A 2-fold increase in precision was observed in both 2D (XY-plane) and 1D (Z-axis) localization, as has previously been noticed in several super-resolution-based methods [13, 14]. Unlike existing methods that require sophisticated PSF engineering or specific materials, here, we have achieved higher photon efficiency in 3D localization with a diffraction-limited point-scanning confocal microscope.

In this work, we investigate the potential of non-uniform illuminaton in the context of real-time 3D single-particle tracking (RT-3D-SPT) [1]. Developed by several groups over the past decades [15–23], RT-3D-SPT uses active feedback to keep a single particle at the center of the microscope objective’s focal volume. In 3D-SMART, the laser spot is guided to sample the XY-plane following a Knight’s Tour pattern and simultaneously sampling along the Z-axis following a sine wave, creating a scanning volume (Fig. 1a-c) that samples the vicinity of the particle of interest in an approximately uniform manner (Fig. 1d). Since the particle is held stationary in the lab frame, *a priori* information about the particle position is available during the measurement. The question then occurs: If *a priori* information is available regarding the particle’s position, can the above-described precision limit be surpassed? We explore this potential in the following work. By examining the expected information extracted from photons collected at various locations relative to the particle center, we show that off-center sampling leads to dramatically increased precision compared to uniform illumination schemes. We then demonstrate that this photon-efficient sampling can be applied in practice using a 3D patterned laser spot. The sampling density of an off-center, information-efficient pattern in 3D is shown in Fig. 1e.

**Figure 1.**
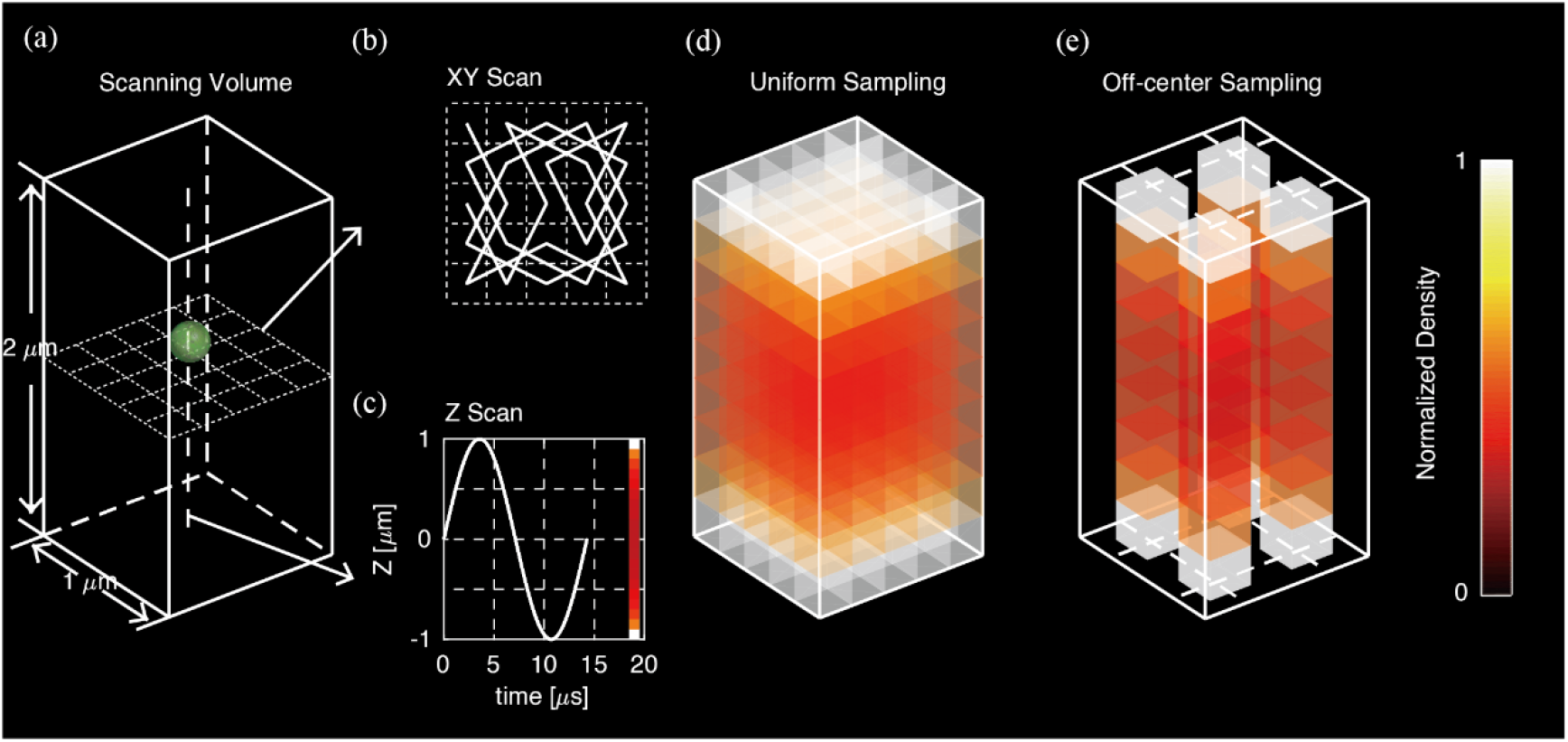
(a) Complete laser scanning volume with a dimension of 1×1×2 μm implemented in 3D-SMART. (b) The laser is scanned in a Knight’s Tour pattern in the XY-plane using an EOD. (c) Along the Z-axis, a TAG lens is used to drive the focus in a sinusoidal pattern. Color bar indicates photon arrival density along the Z-axis. (d) Scanning density in the volume when sampling in the default pattern shown in (b-c). (e) Scanning density of a 3D information-efficient pattern, with the XY-plane, scanned in an off-center, information-efficient pattern. The Z-axis is scanned with a sine wave with the laser power unmodulated.

## Theory

The probability density of observing a photon from an emitter in a microscope in one dimension is approximated by:

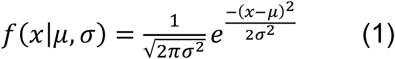

Where *μ* is the particle position and *σ* is related to the width of the PSF. The Gaussian distribution is a good approximation for the actual diffraction pattern of a point-source, which is an Airy function [24]. The size of the PSF is ultimately limited by diffraction and not user adjustable. The prefactor 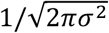 is a normalization factor and will be neglected for the rest of this discussion. The Fisher Information (FI) is a statistical measure employed to quantify the amount of information expected when estimating a parameter of the underlying distribution [25]. The FI (*J*) is inversely related to the precision and is given by:

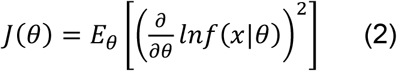

Here *θ* is a parameter (or vector of parameters) of the underlying distribution, and *E_θ_* is the expectation value. The FI for equation (1) above is:

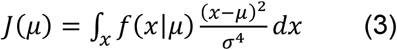

Taken over all space, the expectation value above yields a constant value of 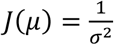, the average amount of FI contributed by each observed photon. The expectation is replaced by the integral in this expression. It is also noticeable that *σ* will be the constant prefactor upon integration and does not affect the solution. For a total of *N* photons, the FI is simply 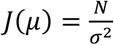. The FI is the inverse of the expected variance, so it is straightforward to see that this is simply the limit of localization precision *(σ/√N)* described above. However, this is only the average value, and photons collected from certain parts of the distribution contribute more to the overall FI than others. To see this, we can more closely examine the integrand above.

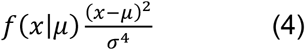

This integrand, which we will refer to as the “information density,” is plotted in Figure 2 to show the contribution of observed photons versus *x − μ*(so the origin is the particle position). From Figure 2, it is easily seen that there are three critical points. There is a minimum in the information density at *x = μ*, and two maxima at 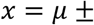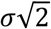. The minimum at *x = μ* contributes zero FI, meaning that photons collected precisely from the particle center yield no information on the particle position. The maxima indicate that photons collected from the off-center positions yield the most FI. In a typical imaging experiment, which employs uniform illumination, the above analysis is not applicable. First, the particle’s location is unknown, so it is impossible to collect photons only from specific areas around the particle’s position. Second, while the photons collected exactly from the particle center yield zero FI, there is no downside to collecting them if unlimited photons are available. However, there are experimental conditions under which it is possible and even desirable to tailor the excitation pattern. For single-molecule tracking or super-resolution imaging, a finite number of photons can be extracted from each molecule before irreversible photobleaching occurs. In these experiments, it is therefore beneficial to collect photons from high information areas only if possible. That being said, this approach is typically impossible because there is no *a priori* information of the particle position. This is where the *a priori* information regarding the particle position available from active-feedback single-molecule tracking comes in.

**Figure 2.**
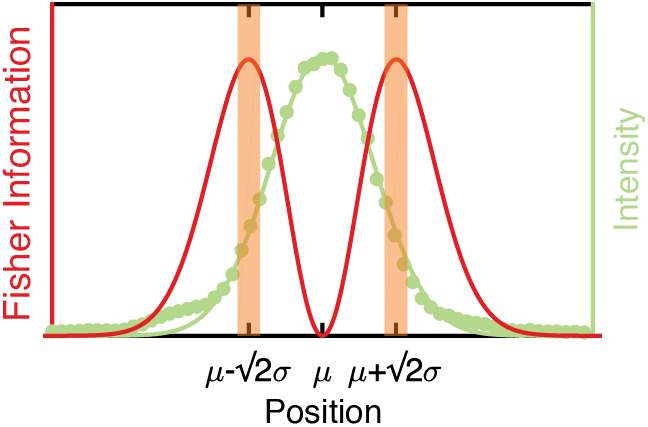
Demonstration of the proposed information-efficient sampling in a single dimension. Green dots represent the experimentally observed intensity of a bead along the X-axis, and the green line shows the Gaussian fit of the PSF. The red line shows the information density of the PSF. Orange shaded areas indicate photons with high information density.

In the following section, we survey three-dimensional laser scanning patterns to identify the most information-efficient sampling patterns. We start with a discussion of applying this sampling along the XY-plane and Z-axis separately, followed by a discussion of achieving isotropic information-efficient sampling in all three dimensions.

## Results

### Identification of a photon-efficient sampling pattern in the XY-plane

In the RT-3D-SPT system reported by Hou et al. [12], a focused laser spot is guided by an electro-optic deflector (EOD) to scan a 5×5 grid with 250 nm between adjacent pixels in the XY-plane (Fig. 3a). Each pixel is sampled for 20 μs in a scanning cycle. The 5×5 grid, typically scanned in a knight’s tour pattern, ensures a uniform illumination pattern. However, the FI-based analysis above suggests that off-center positions should be selectively sampled to achieve the highest photon efficiency. To obtain an unbiased search for photon-efficient patterns, we evaluated subsets of the default 5×5 pattern. The 25 pixels were divided into six different groups of inequivalent pixels based on their distance to the scan center (Fig. 3b). Immobilized fluorescent beads distributed in PBS buffer were then scanned using the default 5×5 EOD pattern in the XY-plane. A piezoelectric stage was then used to step the particle position in 20 nm increments through the center of the scan area. All 62 possible combinations of inequivalent patterns (Fig. S1) were tested to identify the most photon-efficient sampling pattern. In each experiment, photons were collected from each of the 25 pixels, but only photons obtained from pixels in a given pattern were used to perform data analysis.

**Figure 3.**
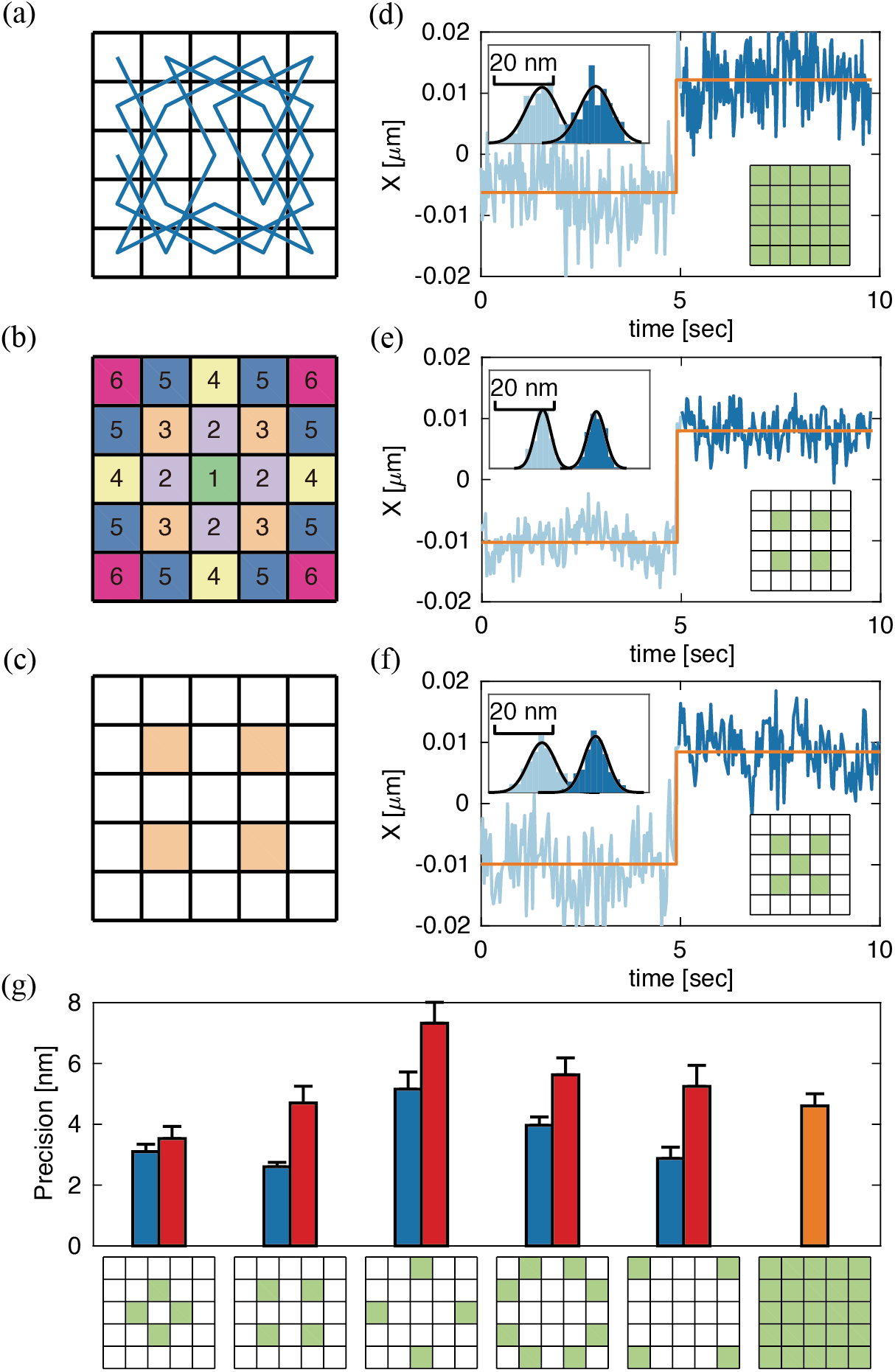
(a) 5×5 Knight’s Tour scanning pattern. (b) Inequivalent pixels based on distance from the scan center. (c) 4-pixel 4-Corners scan, which uses only pixels from pattern 3 in (b), that was found to have the highest precision. (d-f) MLE positions versus stage positions obtained with the default 5×5, 4-Corners, and 4-Corners plus center pixel scan patterns. The average standard deviation of the estimated positions was measured to be 5.5, 2.8, and 4.4 nm, respectively. The orange line shows the relative stage position. The dark and light blue lines show the estimated position calculated for each 2000 photons. The different shades of blue indicate estimation based on two different stage steps, 20 nm apart. (g) Precision obtained from selectively using photons obtained from pixels in different inequivalent patterns (pattern 2-6 in 4b) alone (blue), specific pattern plus center pixel (red), and default 5×5 pattern (orange).

For each different sampling pattern, the particle position was estimated using maximum likelihood estimates (MLEs). In MLE, the likelihood *(L)* for a position estimate *(μ)* from an arbitrary number of photons (*N*) is defined as the product of probability density of photon arrival positions based on a given model *f* which is a function of *μ*:

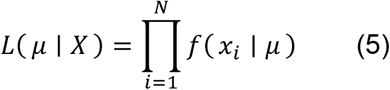

Here the model *f* is a Gaussian distribution described in (1), *X* refers to the set of arrival positions *x_i_(i* = 1,2…*N*) of photons used for estimation. The best estimate for the particle position *μ* is obtained when *L* is maximized. A detailed example of MLE is shown in Fig. S2. Localization precision at different numbers of photons of each estimation is shown in Fig. S3. In this study, every consecutively received 2000 photons are used for each position estimate unless otherwise stated.

MLE analysis was performed on immobilized 190-nm fluorescent beads that were stepped in 20 nm intervals over a 100 nm range for the XY-plane. The average standard deviation of MLE positions at different stage positions was used to quantify the precision (Fig. 3d-f). It was observed that the MLE positions were proportional, but not exactly equal, to the expected particle positions. Off-center sampling generally results in an overestimation of the actual particle motion compared to uniform sampling, which generally yields realistic position estimation. An underestimation of the actual particle motion is associated with sampling patterns in which the center pixel is oversampled compared to uniform sampling. This observation was validated by a simulation of sampling a 2D Gaussian emitter, and results were shown in Fig. S4. Calibration was performed to account for these differences in response to the same changes in particle position (Fig. S5).

Interestingly, though we did not propose an *a priori* model based on Fisher information, the unbiased search resulted in an off-center, FI-efficient pattern. One 4-pixel pattern, which we refer to as the 4-Corners pattern (Fig. 3c), yielded the highest precision of 2.6 ± 0.3 nm, compared with the default pattern (where all 25 pixels are sampled) which gave 4.5 ± 0.3 nm precision (Fig. 3d,e). In both cases, localization was performed using 2000 photons. Additional data were obtained with the laser excitation matching the 4-Corners pattern to validate that selectively using specific photons in data post-processing was equivalent to sampling with the designated pattern. The resulting precision was 2.8 ± 0.4 nm, in excellent agreement with the post-acquisition processed data above. The effect of the number of photons on the relative advantage of the 4-Corners pattern was also investigated, showing a doubling in precision for various values of *N* (Fig. S6). Notably, when the center pixel is added back to the 4-Corners pattern, the precision is nearly two-fold worse than the 4-Corners pattern alone and comparable to the full 25-pixel scan (4.7 ± 1.1 nm vs. 4.6 ± 0.8 nm, Fig. 3f). Complete 2D trajectories of Fig. 3d-f were shown in Fig. S7. We note here that different sampling patterns yield different emission rates for the same laser power. For example, the default 5×5 pattern gives a roughly 1.5-fold increase in intensity compared to the 4-Corners pattern, but the 4-Corners pattern still exhibits doubled-precision when sampling with equal bin time (Fig. S8). A more thorough investigation of patterns consisting of inequivalent pixels No. 2-6 (Fig. 3b) with or without the center pixel showed that precision obtained with the center pixel was always worse than without (Fig. 3g). The poor precision upon sampling the center pixel shows the importance of not sampling low FI areas around the particle.

### Laser modulation in Z-axis

We then proceeded to achieve photon-efficient sampling along the Z-axis. A Tunable Acoustic Gradient (TAG) lens [26, 27] was used to create custom illumination patterns along the axial direction. The TAG lens deflects a focused laser spot in a sine wave with an amplitude of ~1 μm around the focal plane at a frequency of ~70 kHz (Supplementary Figure 4). The following mathematical relation describes the probability density of this sine wave with an amplitude of 1:

**Figure 4.**
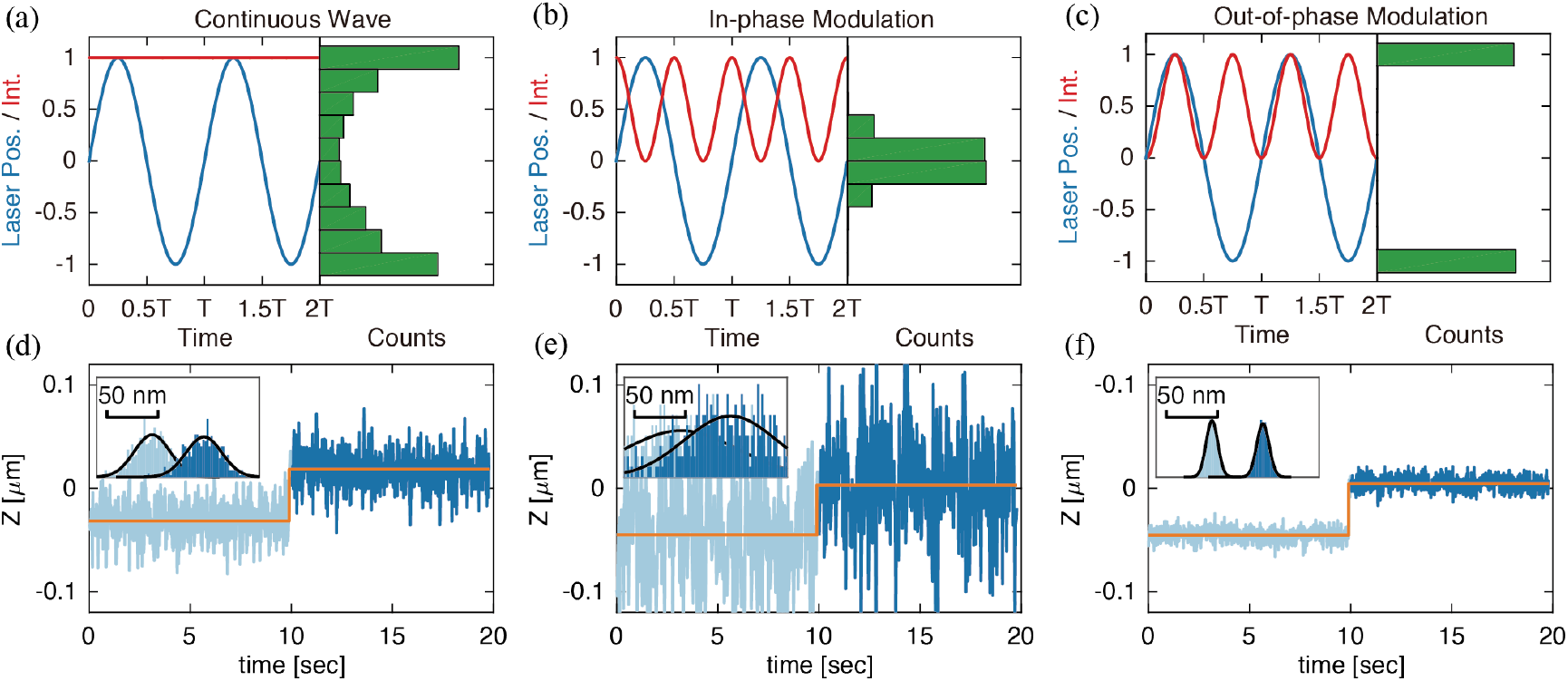
(a-c) Laser intensity versus laser position in Z-axis with photon arrival distribution of 10000 photons from imaging a 190 nm fluorescent bead with TAG lens scanning in unmodulated (continuous wave) or modulated power (no XY scanning). Note that the scanning rate of the TAG lens was held constant at 70 kHz. (d-f) Estimated particle position versus stage position obtained from step tests. Each position estimation was based on 2000 photons. The average standard deviation of estimated positions at each stage position in (d) CW, (e) in-phase modulation, and (f) out-of-phase modulation from the partial trajectories shown in the figures was measured to be 17.6, 50.3, and 5.9 nm, respectively.

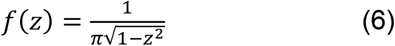

The probability density has a distribution where most probability is piled up at the edges (top and bottom) of the scanning volume (Fig. 1b). Photon arrivals obtained by scanning an immobilized fluorescent bead with a non-modulated, continuous wave (CW) laser spot confirms this distribution (Fig. 4a). According to the theory described above, photons collected at a certain distance away from the center contain the highest information density and lead to the most efficient sampling. However, using CW laser modulation, many photons are still collected from the center of the scanning volume (where the particle spends most of its time). Real-time modulation of the laser intensity was applied to shift the photon arrival distribution away from this low information density area. To do so, the frequency and phase of the TAG lens were captured by a field-programmable gate array (FPGA, NI-7852R). A digital signal with the same frequency and adjustable phase delay was sent to a lock-in amplifier (SR850, Stanford Research Systems) and then to a multiplier circuit to double the original frequency. The frequency-doubled signal was then used to modulate the laser’s power, creating custom illumination patterns along the Z-axis. A detailed illustration of how the signal was processed in the system is shown in Fig. S9. Phase delays between 0o to 90° were tested. Figure 4 shows photon arrival distribution from immobilized 190 nm fluorescent beads sampled at each of the various conditions (CW, in-phase modulation, out-of-phase modulation). At 0o phase delay, photon arrivals occurred at the imaging volume center (in-phase modulation, Fig. 4b). When the delay was 90°, (out-of-phase modulation) photon arrivals were clustered at the edges, with a minimal number of photons at the center (Fig. 4c). Complete trajectories of data shown in Fig. 4d-f were shown in Fig. S10.

Upon performing step tests and data calibration similar to the XY scanning above, the average precision of particles scanned with CW, in-phase modulation, and out-of-phase modulation were 18.2 ± 2.3, 52.0 ± 24.2, and 9.2 ± 1.3 nm, respectively (Fig. 4d-f). These results reaffirmed that off-center sampling (out-of-phase modulation) is the most information-efficient along the Z-axis, as precision was nearly doubled compared with CW power (9.2 vs. 18.2 nm). In-phase modulation, which only samples near the particle center, yielded very poor position estimates.

### Determination of optimized sampling parameters in 3D

Photon-efficient sampling patterns with the ability to localize in all three dimensions were determined by step tests similar to those described above. Step tests with the full 25 XY pixels and CW laser modulation along Z were first performed as a reference. These yielded precisions of 9.3 ± 0.8 nm in X and 22.2 ± 1.6 nm in Z (n = 5). Step tests were then performed with the 4-Corners EOD pattern and CW laser, which gave average precision of 6.8 ± 0.6 nm in X and 19.6 ± 1.7 nm in Z, confirming the advantage of information-efficient off-center sampling. A comparison of step tests in X obtained with the default 5×5 pattern and the 4-Corners EOD pattern with TAG lens operating in CW power is shown in Figure 5. These results again confirm the importance of not collecting photons from the center of the 3D volume. The magnitude of the EOD scale (size of each pixel) and amplitude of TAG lens scan that gave the highest precision in X and Z were found to be 200 nm and 30% TAG lens amplitude (FWHM = 1.93 μm), respectively (Fig. S11). Apart from the amplitude of the TAG lens, the laser power modulation pattern also altered the precision. When imaging with 4-Corners and out-of-phase modulation, the Z precision was high (12.6 ± 0.4 nm), but the X precision was low (22.8 ± 3.1 nm). Inversely, 4-Corners and in-phase modulation resulted in high X precision (4.7 ± 0.2 nm) and low precision in Z (36.0 ± 5.0 nm). Photon arrival distribution along the Z-axis of 190-nm beads sampled with 4-Corners pattern and modulated power at different phase delay is shown in Fig. S12. It is noticeable that out-of-phase modulation in axial-only scanning resulted in a “bi-plane” distribution (Fig. 4c), similar to previously reported work [28]. It should be noted that when XY scanning is enabled, photons from non-edge positions are not necessarily excluded (Fig. S12h).

**Figure 5.**
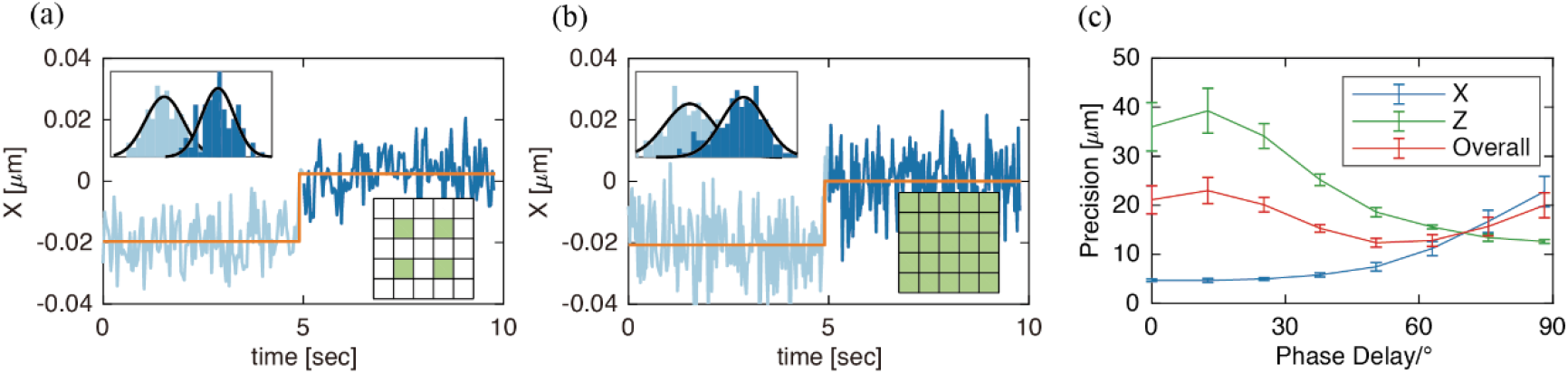
(a-b) Estimated position versus stage position sampled with the 4-Corners and default patterns. Both used CW laser power modulation along the axial direction. The standard deviation of estimated positions (based on 2000 photons) was (a) 7.4 and (b) 10.5 nm from the partial trajectories shown above. (c) X, Z, and 3D precision of 5 different immobilized 190-nm fluorescent beads sampled with 4-Corners in the XY-plane and modulated power at different phase delay along the Z-axis.

To get a comprehensive standard of overall precision in all dimensions in each condition, we define an overall 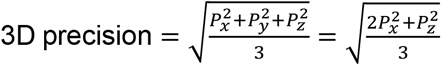. The 3D precision is derived from X and Z precision here since X and Y were sampled via the same mechanism. Step test trajectories obtained with the 4-Corners and modulated laser power with a phase delay of 50.4° yielded the highest 3D precision (12.3 ± 0.9 nm). This result is comparable to the 3D precision found for the 4-Corners pattern and CW laser power (12.6 ± 1.1 nm). It is also noteworthy that it is possible to achieve equivalent X and Z precision when a phase delay of ~70° is applied, as is shown in Fig. 5c, where the X and Z precisions intersect. The estimated precision at this point is *P = P_x_ = P_z_* = 14.3 *nm*. This condition makes it possible to conduct isotropic sampling despite having non-uniform point spread function scales in different dimensions (Fig. S13c-d).

### Discussion and Conclusion

This study showed that implementing information-efficient laser scanning patterns led to dramatic improvements in precision in all three dimensions. A 4-pixel, 4-Corners pattern yielded the highest precision of 2.6 ± 0.3 nm in the XY-plane compared to 4.6 ± 0.8 nm given by a default 5×5 pattern (43.5% increase). When sampling the Z-axis only, out-of-phase modulation of the laser power relative to the TAG lens phase (which gave a bi-modal distribution of photon arrivals) gave the highest precision of 9.2 ± 2.6 nm compared to 18.2 ± 4.6 nm given by CW power modulation (49.5% increase in precision). In 3D scanning, the 4-Corners pattern with laser power modulated with a 50.3° phase delay gave a 3D precision of 12.3 ± 0.9 nm compared to 14.9 ± 1.1 nm given by the default 5×5 pattern with CW power modulation (17.5% increase). These results are consistent with the hypothesis that sampling at higher Fisher information regions leads to the best precision. Moreover, it is also shown that sampling directly at the center of the particle is inefficient, with those photons carrying little or no information. Achieving high photon efficiency is extremely important as typical fluorophores give out only a limited number of photons. The off-center, information-efficient imaging proposed in this work is the first to achieve higher photon efficiency by merely scanning a focused laser spot without requiring any PSF engineering.

It is noticeable that RT-3D-SPT methods might have used similar off-center excitation patterns. For example, orbital tracking methods developed by Gratton [29], Mabuchi [30, 31], Lamb [20, 32], and others utilize a circular scanning pattern in the XY-plane, which minimizes sampling of the particle center. The motivation for such an approach was to sense changes in the particle position by modulating the particle’s intensity. Others have utilized a method where the laser is scanned across the vertices of a tetrahedron [33, 34]. All of these methods benefit from the off-center sampling that we demonstrate above. Unlike previous works[15, 21] that utilized scan-free localization, using the EOD and TAG lens here is advantageous in defining a custom excitation pattern due to their highly-tunable nature and the intrinsically information-efficient pattern caused by the TAG lens’s sinusoidal motion.

Our work revealed the importance of information-efficient excitation patterns in particle localization and tracking using non-uniform illumination. The information-efficient patterns investigated in this work could shed light on an emerging field in biology: slowly moving particles. Recent studies have shown that even the previously considered “static” intracellular vesicles could still undergo small-scale motions of different rates, which could profoundly influence the fate of these particles [35]. Information-efficient sampling should optimize combined spatiotemporal precision in such demanding experiments, where the diffusive step sizes are extremely small and the fluorophores short-lived.

