## Supplementary Figures and Tables for "Information-Efficient, Off-Center Sampling Results in Improved Precision in 3D Single Particle Tracking Microscopy"

**Supplementary Information**

**Table of Contents**


| **Section** | **Page** |
|---|---|
| Possible Excitation Patterns in a 5×5 Grid | 2 |
| Maximum Likelihood Estimation | 3 |
| Localization Precision at Different Number of Photons | 4 |
| Simulation of Influence of Different Sampling Patterns on Measured Position versus Actual Position | 5 |
| Calibration of Step Test Data | 6 |
| Precision from Different Number of Photons | 9 |
| Sample 2D Trajectories | 10 |
| Precision from Equivalent Sampling Intervals | 11 |
| Laser Modulation along the Z-Axis | 13 |
| Sample Z Trajectories | 14 |
| Optimized EOD and TAG Lens Scales | 15 |
| Photon Arrival Distribution from Different Phase Delays in 3D Sampling | 16 |
| Measurement of the Point Spread Function | 17 |
| Precision under Different Experimental Conditions | 18 |
| Schematic of 3D-SMART setup | 19 |

**Possible Excitation Patterns in a 5×5 Grid**

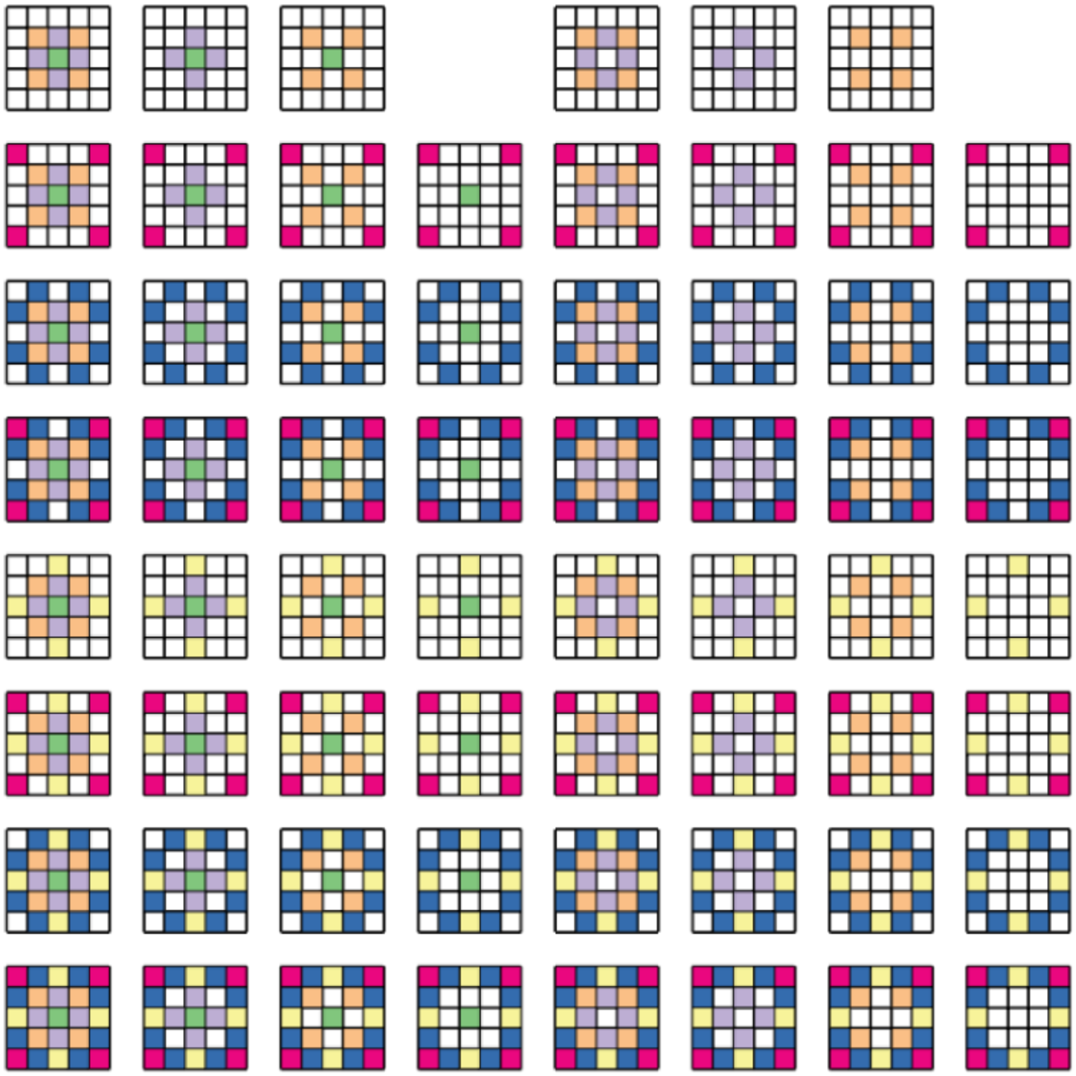

**Supplementary Figure 1.** All 62 possible excitation patterns from random combinations of 6 inequivalent pixels shown in Fig. 3b.

**Maximum Likelihood Estimation**

To estimate the position $\mu$ of an emitter, consider a Gaussian distribution $f(x \mid \mu)$ and let $X$ be a series of photons collected at different laser scan positions $x_i$ $(i = 1,2, \dots, N)$ to be used to give one estimation:

$$f(x|\mu) = \frac{1}{\sqrt{2\pi\sigma^2}} e^{\frac{-(x-\mu)^2}{2\sigma^2}} \quad (1)$$

The likelihood of a particular position $\mu$ given the observed photons $X$ is given by:

$$L(\mu \mid X) = \prod_{i=1}^{N} \frac{1}{\sqrt{2\pi\sigma^2}} e^{-\frac{(x_i-\mu)^2}{2\sigma^2}} \quad (2)$$

To maximize the likelihood given by the likelihood product, we take the natural log of the above expression:

$$logL(\mu|X) = -nlog(2\pi\sigma^2) - \sum_{i=1}^{N} \frac{(x_i - \mu)^2}{2\sigma^2} \quad (3)$$

The first derivative of $logL(X \mid \mu)$ with respect to $\mu$ is then used to obtain the value of $\mu$ that maximizes $L$. The problem can then be simplified to find a $\mu$ that satisfies:

$$\frac{\partial L(\mu|X)}{\partial \mu} = \sum_{i=1}^{N} \frac{x_i - \mu}{\sigma^2} = 0 \quad (4)$$

The solution is:

$$\mu = \frac{1}{N} \sum_{i=1}^{N} x_i \quad (5)$$

The result reveals that when the underlying distribution is Gaussian, the maximum likelihood estimate of particle position $\mu$ is the mean of photon arrival positions. An example of MLE of X position is shown in Fig. S2.

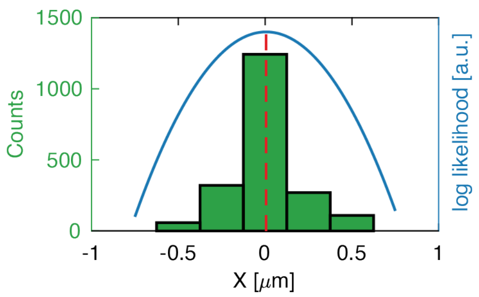


**Supplementary Figure 2.** Maximum Likelihood Estimation of X position. Green bars show the distribution of 2000 consecutively collected from sampling a fixed 190-nm bead with the default EOD pattern (all 25 pixels, TAG lens off) in a 34 ms interval. The blue curve shows the log-likelihood of different estimations based on positions of the photons collected, and the estimated particle position with the highest likelihood was 6 nm (dotted red line, rounded to nearest nm).

**Localization Precision at Different Number of Photons**

When maximum likelihood estimation is used to obtain particle position, it is possible to choose an arbitrary number of photons for each estimation. Fig. S3 shows the average particle localization precision versus the number of photons along with the theoretical lower limit for each estimation obtained from scanning immobilized 190-nm fluorescent beads. The difference between the X and Z precisions results from the larger size of the PSF along the Z-axis. It should be noted that the Z precision does not actually break the theoretical lower bound, as shown in Fig. S5, since the data are not calibrated to reflect actual particle motion.

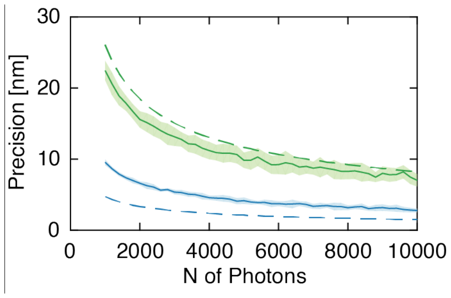


**Supplementary Figure 3**. Localization precision versus the number of photons used for analysis in X (blue curve) and Z (green curve). Dashed lines indicate theoretical lower bound determined by $\sigma/\sqrt{N}$. Each data point was from 9 different particles. The standard deviation of data points is shown by shaded areas.

**Simulation of Influence of Different Sampling Patterns on Measured Position versus Actual Position**

A simulation was performed to investigate the cause of underestimation or overestimation of the actual position when different sampling patterns were applied. Here, we modelled an emitter that follows 2D Gaussian distribution where intensity $I$ at a given point $(x, y)$ in a 2D plane is given by:

$$I(x, y) = I_0 e^{-\frac{(x-x_0)^2+(y-y_0)^2}{2\sigma^2}} \tag{5}$$

Here $(x_0, y_0)$ is the emitter position, $I_0$ is center intensity and $\sigma$ is the uncertainty of the Gaussian point. A custom grid containing $N$ pixels that is equivalent to any pattern is built and intensity at each pixel can be easily obtained by substituting center coordinates $(x_k, y_k)(k = 1,2 \dots N)$ of each pixel into (5):

$$I_k = I(x_k, y_k) \tag{6}$$

Photon arrival probability $P_k$ at each pixel can then be approximated as normalized intensity:

$$P_k = \frac{I_k}{\sum_{k=1}^{N} I_k} \tag{7}$$

Simulated photon arrival data were then obtained through a simple discrete probability distribution model and analyzed. Uncalibrated simulated step test data are shown in Fig. S4. As is seen in Fig. S4, the simulated results agree with the actual experimental data, where even sampling (Fig.S4a) gave accurate estimation as expected. In contrast, the “off-center” 4-Corners pattern (Fig. S4b) overestimated the real position. Adding a fifth center pixel resulted in underestimation (Fig. S4c).

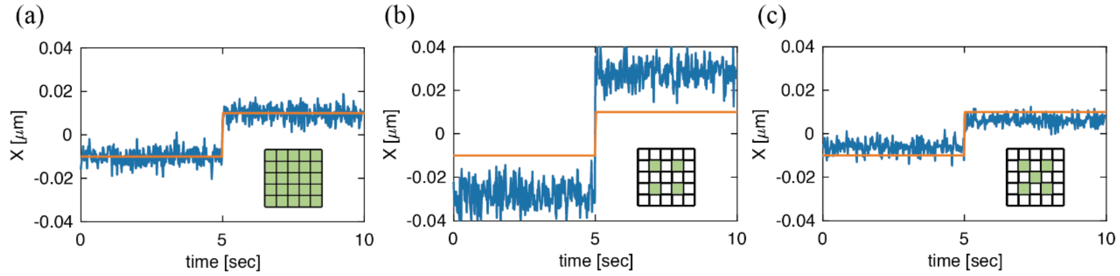


**Supplementary Figure 4.** Simulated step test data with laser scanning in (a) default 5×5 pattern, (b) the 4-Corners off-center pattern, and (c) the 4-Corners plus center pixel pattern. The simulated particle was moved in 20 nm steps in all simulations. The measured steps from simulation were (a) 19.5 nm, (b) 56.1 nm, and (c) 12.1 nm.

**Calibration of Step Test Data**

Immobilized 190-nm fluorescent beads were stepped through the laser scanning volume using a piezo stage to evaluate the fidelity of MLE. Estimated positions were then compared with the known stage motion to calibrate positions given by MLE for different scan patterns. A step test of a 190-nm fluorescent bead imaged with the default EOD pattern, and no axial scanning is shown in Fig. S5a. In the trajectory, the stage moved 20 nm every 10 seconds along the X-axis. The estimated positions display stepwise motion over a considerable range (>150 nm). Stage positions were aligned with a given step in which the average of MLE positions was closest to 0. Notably, the MLE positions from particles sampled with the default EOD pattern were close to the set stage positions, showing an average slope of 1.2 ± 0.1. The slope was calculated over a 100-nm range around the zero position (Fig. S5b).

It was noticed that relative motion of estimated position versus actual position varied significantly for other sampling patterns. For example, the slope between the MLE positions and the applied stage position was 2.2 ± 0.2 for the 4-Corners EOD pattern. When axial scanning with CW power was applied, this slope decreased to 0.71 ± 0.04. Though different sampling patterns yielded different slopes, it was simple to restore the fidelity of the MLEs by applying the following equation:

$$x_{real} = x_{est}/k \quad (8)$$

Here $x_{est}$ is the position given by MLE, $x_{real}$ is the actual position calibrated to account for the linear correlation between the estimated position and the actual position. The calibration factor $k$ is the slope obtained from a linear fit of stage positions and MLE positions for a given sampling condition. This simple calibration method makes it immediately clear that when uncalibrated MLE positions display comparable precision for different scan patterns, a higher value for $k$ results in higher precision.

For example, a trajectory obtained from a step test of a 190-nm fluorescent bead sampled with the 4-Corners EOD pattern (no axial scanning) is shown in Fig. S5c. The average error of MLE positions (which we refer to as the uncalibrated precision) over five steps was 7.1 nm. This value is worse than the uncalibrated precision for the default 5x5 pattern, which is 5.5 ± 0.2 nm. The 4-Corners pattern, however, gave a very high slope ($k = 2.2$ in trajectory shown in Fig. S5c). A linear fit of the center of MLE positions and stage positions is shown in Fig. S5d. As is shown in Fig. S5e, the error in the estimated position in the highlighted area was substantially smaller once the MLE positions were correctly aligned with the known stage positions, giving a trajectory-wide calibrated precision of 3.3 nm. The uncalibrated precision, calibrated precision, and slope for the default 5x5 pattern and the 4-Corners pattern (no axial scanning) are shown in Supplementary Table S5. Upon calibration, the 4-Corners pattern gave a 43% improvement in precision (2.6 versus 4.6 nm).

For all step test analyses, X step tests were analyzed over a 100 nm range with 20 nm steps. Z step tests were analyzed over a 150 nm range with 30 nm steps. This calibration ensured that MLE positions display a robust linear correlation ( $r^2 > 0.99$) to stage motion.

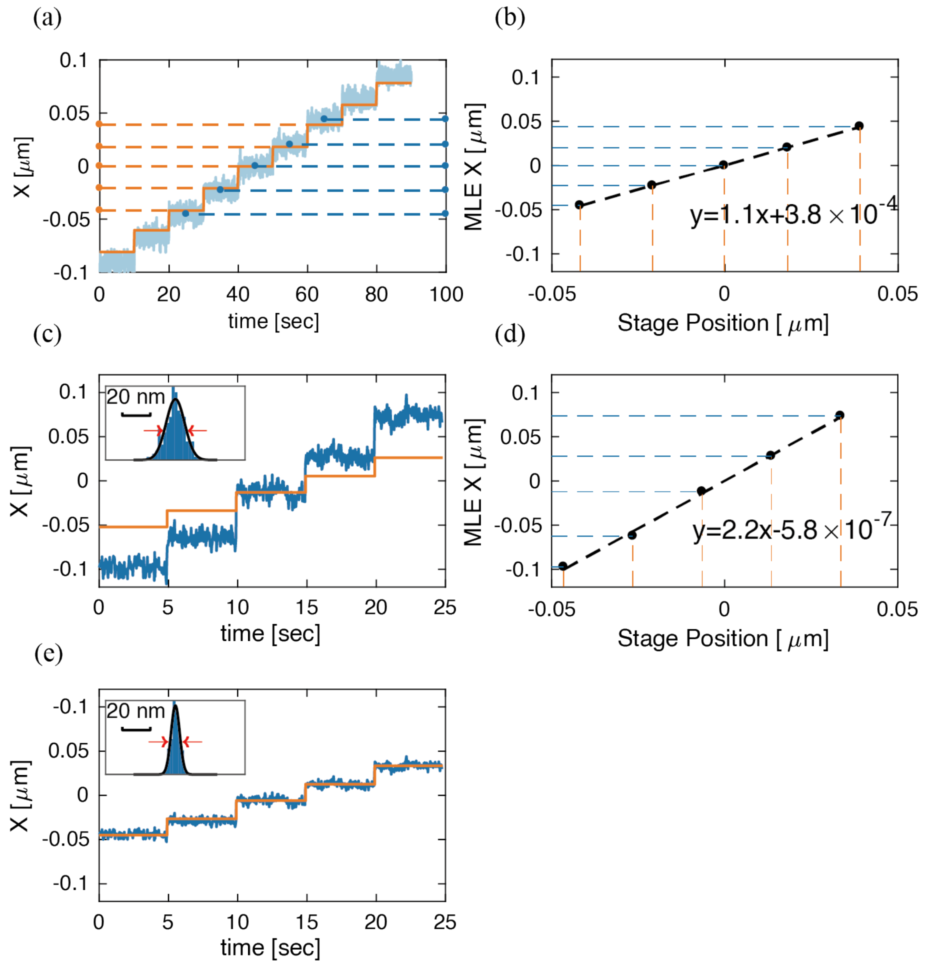


**Supplementary Figure 5.** (a) MLE positions (light blue) versus stage positions (orange) for a 190-nm bead with the default EOD pattern and no axial scanning. The stage was moved along the X-axis in 20-nm steps and stayed at each position for 10 seconds. Stage positions were aligned to MLE positions between 40 and 50 seconds, where the particle was near the center of the scan area. The dashed lines show stage positions (orange) and MLE positions (blue) for each 10-second interval. (b) Linear fit of stage residence positions and center of MLE positions in (a). (c) MLE positions (blue) versus stage positions (orange) for the 4-Corners EOD pattern with no axial scanning. Stage was guided to move 20 nm along X-axis every 5 seconds. Upper left inset shows the distribution of MLE positions between 10 and 15 seconds ($\sigma = 6.7\ nm$). (d) Linear fit of stage residence positions and center of MLE positions in (c). (e) MLE positions from (c) calibrated with a slope of 2.2. Upper left inset shows the distribution of MLE positions between 10 and 15 seconds ($\sigma = 3.1\ nm$).

| Experimental Condition | Uncalibrated Precision/nm | Calibration Factor | Calibrated Precision/nm |
|---|---|---|---|
| Default 5x5 EOD pattern, TAG lens off (n = 11 trajectories) | 5.5 ± 0.4 | 1.2 ± 0.1 | 4.6 ± 0.8 |
| 4-Corners EOD pattern, TAG lens off (n = 5 trajectories) | 5.5 ± 0.2 | 2.2 ± 0.2 | 2.6 ± 0.3 |

**Supplementary Table 5.** Uncalibrated precision, calibration factor, and calibrated precision for two laser scan patterns in XY. The calibration factor was calculated from the average slope over a 100 nm range around the center of the laser scan.

**Precision from Different Number of Photons**

To verify that the improved precision is not specific to a certain number of photons, data analysis was performed with different number of photons. It was observed that the 4-Corners pattern gave a better precision compared to the default pattern or the 5-pixel pattern (4-Corners plus center pixel). The precision at different number of photons per localization from step test data for the 4-Corners pattern and the default 5x5 pattern (no axial scanning) is shown in Fig. S6. Also shown is the theoretical lower limit of $\sigma/\sqrt{N}$ . Here $\sigma$ is 100 nm, which is from the experimentally determined PSF. It should be noted that the theoretical lower limit does not account for the pixel size, background photons, or any other noise sources, which is why it does not overlap with the default 5x5 pattern.

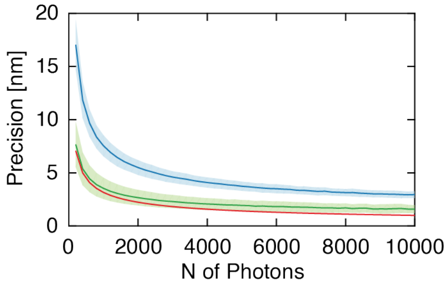


**Supplementary Figure 6.** Localization precision using 200-10000 photons with the 4-Corners pattern (green line) and the default 5x5 pattern (blue line). The red line shows the theoretical lower limit of $\sigma/\sqrt{N}$. Shaded areas indicate the standard deviation from 11 different particles.

**Sample 2D Trajectories**

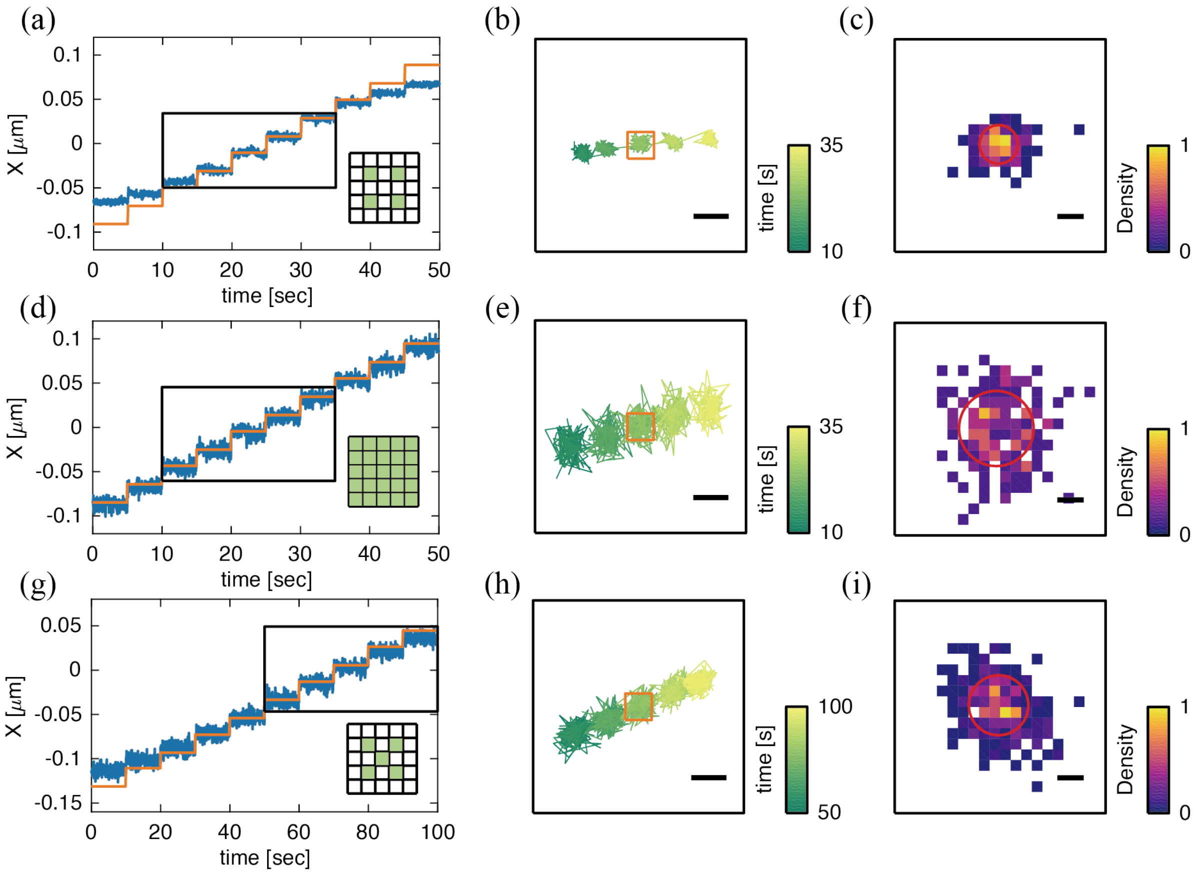


**Supplementary Figure 7.** (a,d,g) MLE positions versus stage positions for three different scan patterns on immobilized fluorescent beads. The stage was guided along the X direction in 10 consecutive steps of 20 nm each. Data are shown for the (a) 4-Corners pattern, (d) default 5×5 pattern, and (g) 4-Corners pattern plus center pixel. The highlighted rectangular areas indicate positions used for calculation of calibration factor and precision. (b,e,h) 2D trajectories of highlighted regions in (a), (d), and (g), respectively. (c,f,g) Density maps of 2D coordinates in highlighted areas in (b), (e), and (h), respectively. Red circles indicate the standard deviation from 2D Gaussian fitting, which yielded (c) 2.66 nm, (f) 5.79 nm, and (i) 4.59 nm. Scale bars: (b,e,h) 20 nm and (c,f,i) 5 nm.

### Precision from Equivalent Sampling Intervals

It is noticeable that different particles have different emission rates, and different sampling patterns result in different intensity at different pixel locations. When using a constant number of photons per localization (N=2000 typically), this can lead to different temporal sampling rates. This difference in the temporal sampling rates does not have any influence on the conclusions drawn with regards to photon efficiency. Fluorescent beads have very high signal-to-background ratio (>100:1) at typical emission rates (50~100 kHz), but it would be a waste of the limited photons to increase the photon flux from a single fluorophore. However, it is still interesting to investigate precision at an equal sampling interval at the same excitation power. Complete X trajectories of 190-nm fluorescent beads sampled with default and 4-Corners EOD pattern and no axial scanning are shown in Fig. S8 (a) and (b). Trajectories in Fig. S8 (a) and (b) were obtained with the same laser power. Each position estimate was from photons collected in a consecutive 20-ms interval. We define the average intensity when the estimated position is closest to 0 as the "center intensity" as an indicator of emission rate of different excitation patterns. The center intensity was measured to be 76.1 and 44.6 kHz for the default 5x5 and 4-Corners patterns, respectively (Fig. S8a,b). Though the default 5×5 pattern gave higher center intensity and thus higher photon count for each estimation, precision was measured to be 6.82 and 3.35 nm in Fig. S8 (a) and (b), which means the 4-Corners pattern still gave higher precision at an even temporal sampling rate.

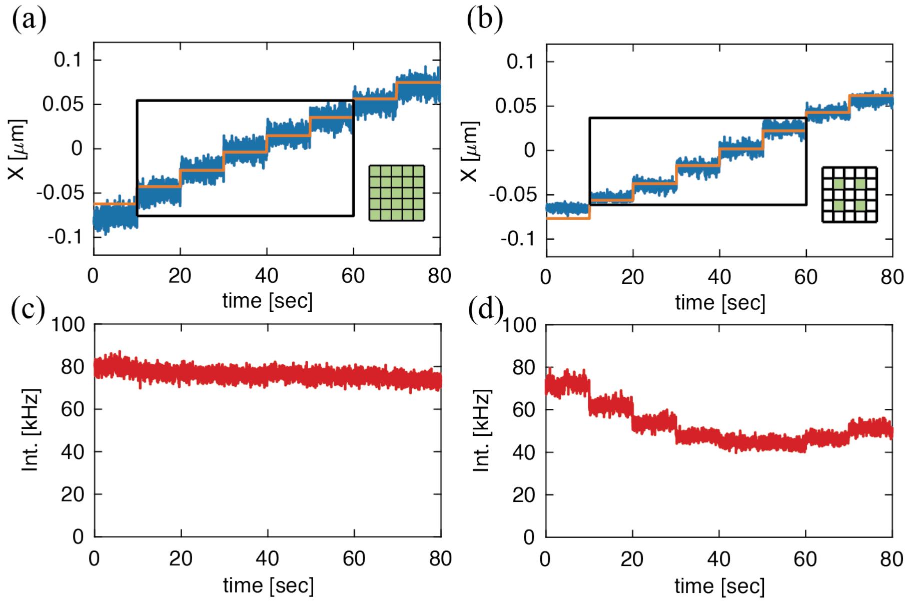


**Supplementary Figure 8.** (a,b) MLE positions (blue lines) vs. stage positions (orange lines) from immobilized 190-nm fluorescent beads sampled with different EOD patterns and no axial scanning. Each position estimate is from photons received in a 20-ms interval. (c,d) Real-time emission rate of trajectories shown in (a,b).

| Experimental Condition | Center Intensity/kHz | Precision/nm | Average N of Photons/estimate |
|---|---|---|---|
| Default 5×5 EOD pattern, TAG lens off | 63.0 ± 2.5 | 7.0 ± 1.4 | 1260 ± 50 |
| 4-Corners pattern, TAG lens off | 44.0 ± 0.6 | 3.4 ± 1.1 | 880 ± 12 |

**Supplementary Table 8.** Center intensity, X precision, and the average number of photons per estimate when the particle is sampled every 20 ms. Each data point is from 11 trajectories.

**Laser Modulation along the Z-Axis**

Laser modulation was achieved through a series of hardware implementations, including an FPGA, lock-in amplifier (LIA), and a frequency-doubling circuit. First, the phase of the TAG lens was captured in real-time on the FPGA. A digital square wave with the same frequency and custom phase delay was then generated (Figure S9a) and sent to the LIA. A sine wave of the same frequency and phase as the square wave was then generated (Fig. S9b) by the LIA and sent to the frequency-doubling circuit. The circuit utilized an AD835 (Analog Devices) 4-quadrant multiplier to multiply two input signals. When identical sine waves (obtained from the LIA) were sent into the two inputs of the multiplier circuit, a signal of doubled frequency was obtained (Fig. S9c):

$$sin^2(2\pi f t + \varphi) = \frac{1}{2} - \frac{1}{2}cos(2\pi(2f)t + 2\varphi) \tag{6}$$

This signal could then be used to modulate laser intensity in real-time.

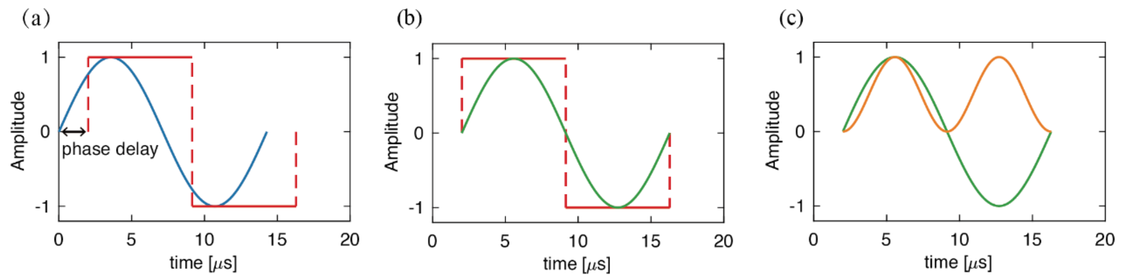


**Supplementary Figure 9.** (a) Original TAG lens control signal (blue) and square wave generated by the FPGA (red). The phase delay could be tuned on-demand for FPGA-generated square wave. (b) Square wave (red) shown in (a) and sine wave generated by the LIA (green). (c) Delayed TAG lens control signal (green) obtained in (b) and laser modulation signal obtained from the frequency-doubling circuit (orange).

**Sample Z Trajectories**

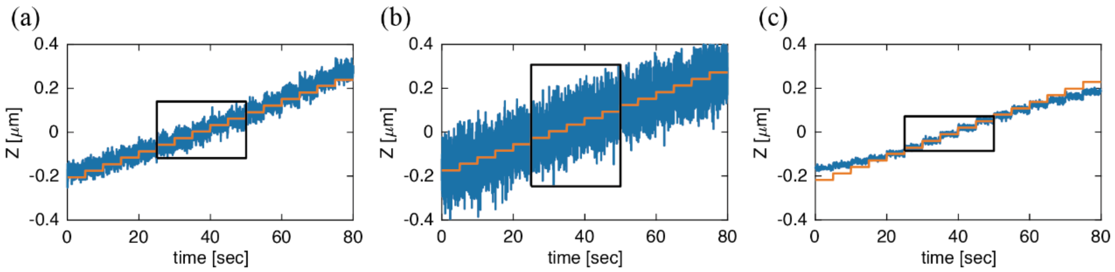


**Supplementary Figure 10**. MLE positions versus stage for three different scan patterns on immobilized fluorescent beads. The stage was guided along the Z direction in 16 consecutive steps of 30 nm each other. Data are shown for the TAG lens working in (a) CW power, (b) in-phase modulation, and (d) out-of-phase modulation (c). Highlighted rectangular areas indicate positions used for calculation of calibration factor and precision.

### Optimized EOD and TAG Lens Scales

Step tests with the 4-Corners EOD pattern and axial scanning with CW laser power were performed using different EOD and TAG lens scales to determine the optimized scanning parameters for 3D localization. It was observed that a 200-nm EOD scale (Fig. S11a) and 30% maximum TAG lens amplitude (Fig. S11b) were independently determined to be optimal for high precision localization. Step tests were also performed to find optimized sampling parameters in single dimensions. When performing XY localization with the 4-Corners pattern (no axial scanning), a 200-nm EOD scale was found to give the highest precision, as is shown in Fig. S11c. It is noticeable that 250-325 nm EOD scale gave an almost identical performance. However, larger EOD scales enabled localization over larger XY ranges, so 300 nm was chosen for further experiments. As is shown in Fig. S11d, when performing localization along Z-axis with TAG lens working in CW power (no XY scanning), 15% amplitude gave the highest precision. Lower amplitude was not tested because TAG lens performance was unpredictable at <15% amplitude, as is shown in Fig. S11b.

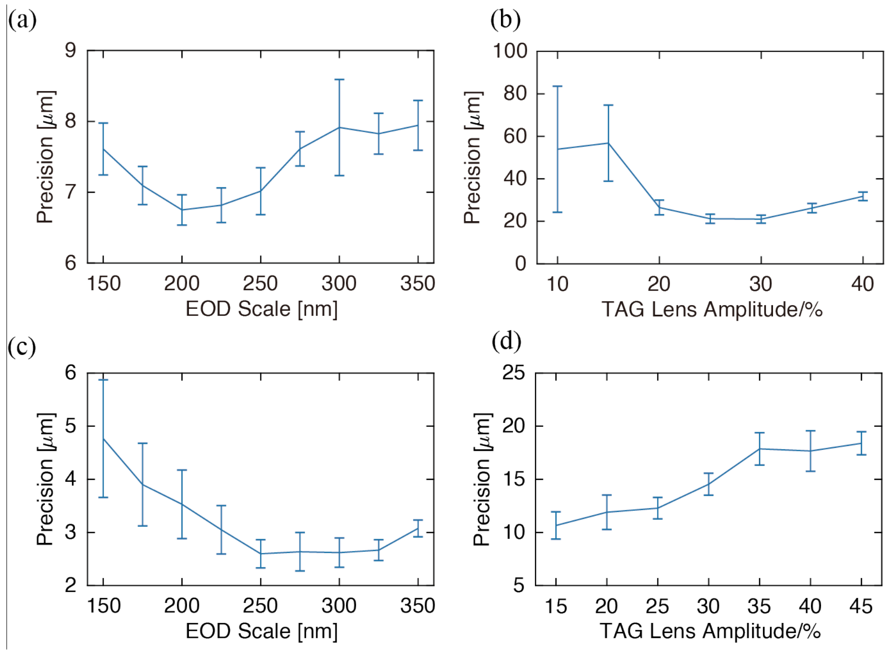


**Supplementary Figure 11**. (a) X precision versus EOD scan size for the 4-Corners pattern with no axial scanning. An optimized precision in X of 6.7 ± 0.2 nm was obtained for 200-nm EOD scale. (b) Z precision versus TAG lens amplitude with CW laser power modulation and 4-Corners pattern at 250-nm scale. An optimized Z precision of 21.0 ± 1.9 nm was obtained at 30% of maximum amplitude. (c) X precision versus EOD scale for the 4-Corners pattern with no axial scanning. An optimized X precision of 2.6 ± 0.3 nm was obtained at 250-nm EOD scale. (d) Z precision versus TAG lens amplitude with CW power and no EOD scanning. All data points were based on data collected from 5 different beads. Each MLE position was calculated from 2000 photons.

**Photon Arrival Distribution from Different Phase Delays in 3D Sampling**

The distribution of photon arrival positions of particles sampled with TAG lens working under different laser power modulation is shown in Fig. S12. These data were taken with the EOD scanning in the 4-Corners pattern. These are slightly different from the data shown in Fig. 4, collected with no XY scanning.

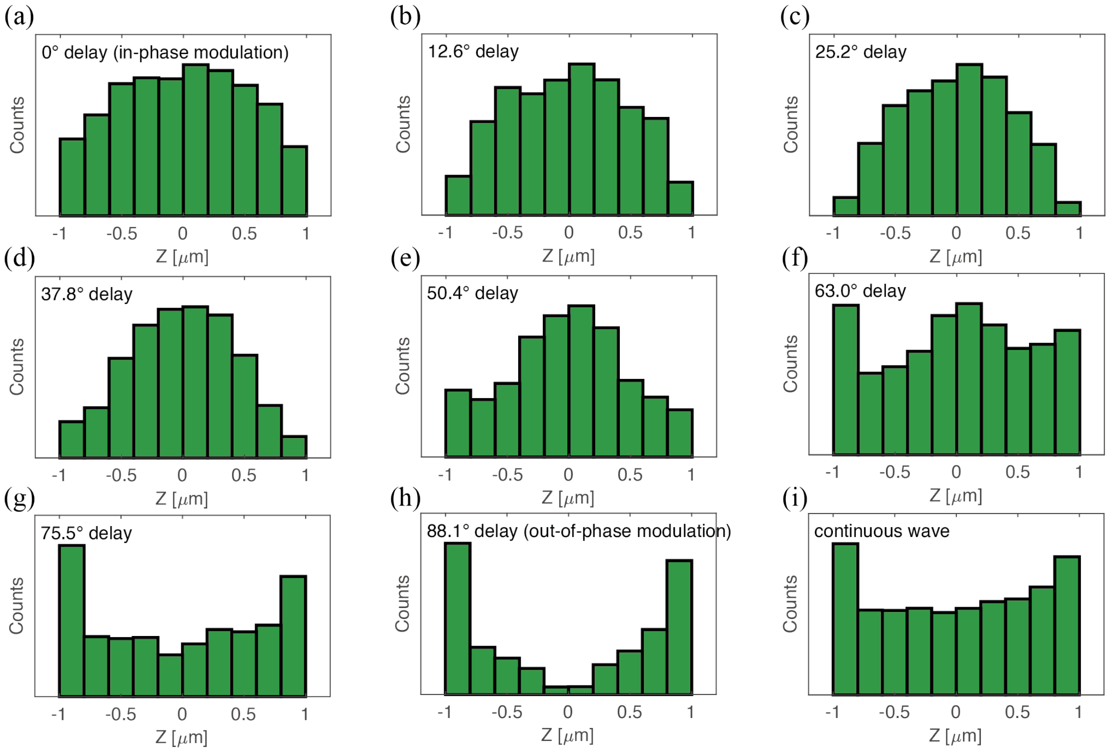


**Supplementary Figure 12**. Photon arrival distribution of fixed 190-nm beads sampled with EOD scanning in the 4-Corners pattern with 250-nm scale and TAG lens working in modulated power at (a-h) different phase delays and (i) CW power. Each figure shows the distribution of 20000 consecutively collected photons from a particle localized near the center of the scanning area.

**Measurement of the Point Spread Function**

To measure the point spread function (PSF), a 190-nm bead fixed on coverslip was scanned point-by-point through a 3D volume of 2 μm x 2 μm x 8 μm, as is shown in Fig. S13a (TAG lens or EOD both turned off). The scanning step in X and Y was 100 nm, and the step in Z was 200 nm (20 out of 41 frames shown for clearer visualization). Intensity at each point was from the integration of photons collected over a 10 ms interval. Interestingly, the PSF scale along the Z-axis was roughly 7-times larger than that in X. With CW laser modulation, the Z precision was only about ~2-3 times worse than the X precision. As shown in Fig. 4a, the TAG lens scan with CW laser power modulation shows a pileup of photon arrivals at the edge of the distribution, which is intrinsically more information efficient than uniform illumination. To show the impact of the TAG lens on the PSF, an image of the same particle in Fig. S13a scanned with TAG lens working in CW power is shown in Fig. S13f, where the size of the PSF is slightly larger (128 nm TAG lens on vs. 99 nm TAG lens off).

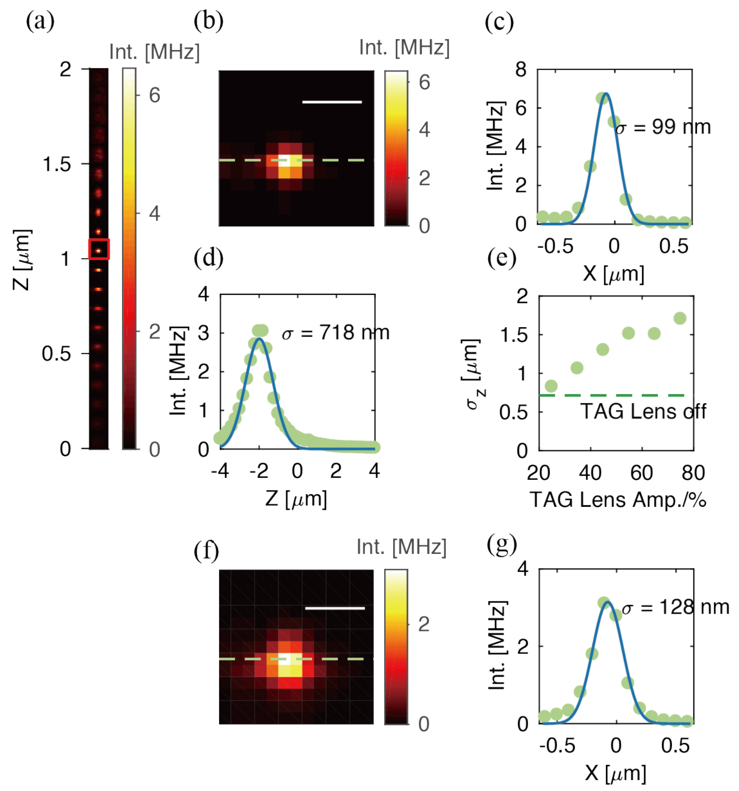


**Supplementary Figure 13.** (a) Scanning image of a 190-nm fluorescent bead. (b) Frame labeled in (a) that gave the tightest focal spot in the XY-plane. (c) Gaussian fit of intensity along X labeled by the green line in (b). (d) Gaussian fit of the intensity of 3x3 pixel area in the center of each frame along the Z-axis as a function of Z. (e) Estimated PSF size in Z at different TAG amplitudes. (f) A frame that gives the tightest focal spot from the same particle shown in (a) scanned with TAG lens working at 25% amplitude. (g) Gaussian fit of intensity along X labeled by the green line in (f). Scale bars in (b) and (f): 500 nm.

**Precision under Different Experimental Conditions**

| Experimental Condition | X Precision/nm | Z Precision/nm | 3D Precision/nm |
|---|---|---|---|
| Default 5×5 EOD pattern, TAG lens off | 4.6 ± 0.8 | | |
| 4-Corners pattern, TAG lens off | 2.6 ± 0.3 | | |
| 4-Corners pattern + center pixel, TAG lens off | 4.7 ± 1.1 | | |
| CW power, EOD off | | 18.2 ± 4.6 | |
| Out-of-phase modulation, EOD off | | 9.2 ± 2.6 | |
| In-phase modulation, EOD off | | 52.0 ± 24.2 | |
| Default 5×5 EOD pattern, CW power | 9.3 ± 0.8 | 22.2 ± 1.6 | 14.9 ± 1.1 |
| 4-Corners pattern, CW power | 6.8 ± 0.6 | 19.6 ± 1.7 | 12.6 ± 1.1 |
| 4-Corners pattern, 50° phase delay | 7.5 ± 0.8 | 18.6 ± 0.9 | 12.3 ± 0.9 |

**Supplementary Table 14.** X, Z, and 3D precision (for 3D sampling patterns) from different experimental conditions

**Schematic of 3D-SMART setup**

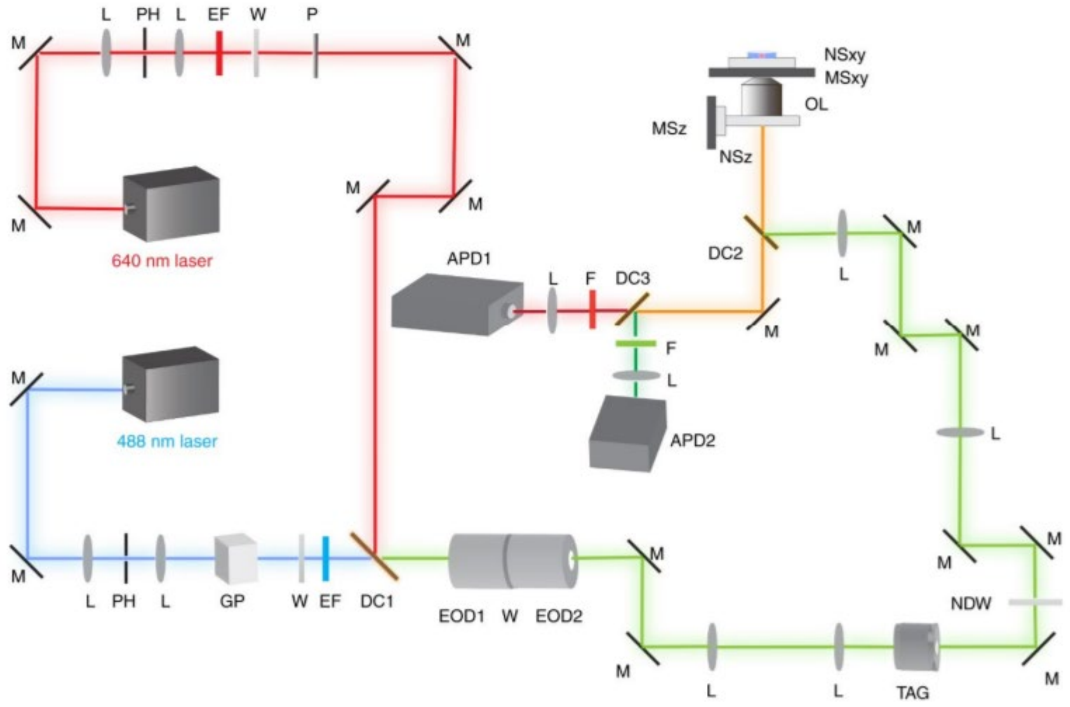


**Supplementary Figure 15.** Schematic of 3D-SMART setup. L: lens; M: mirror; PH: pinhole; GP: Glan-Thompson polarizer; P: polarizer; W: half waveplate; EF: excitation filter; DC: dichroic mirror; EOD: electro-optic deflector; TAG: TAG lens; NDW: Neutral density filter wheel; OL: objective lens; MSxy: xy microstage; MSz: z microstage; NSxy: xy nanopositioner stage; NSz: z nanopositioner stage; F: fluorescence filter. (Figure reprinted from Hou S, Exell J, Welsher K. Real-time 3D single molecule tracking. Nature communications, 2020, 11(1): 1-10.)
